# Diallel panel reveals a significant impact of low-frequency genetic variants on gene expression variation in yeast

**DOI:** 10.1101/2023.07.21.550015

**Authors:** Andreas Tsouris, Gauthier Brach, Anne Friedrich, Jing Hou, Joseph Schacherer

## Abstract

Unraveling the genetic sources of gene expression variation is essential to better understand the origins of phenotypic diversity in natural populations. Genome-wide association studies identified thousands of variants involved in gene expression variation, however, variants detected only explain part of the heritability. In fact, variants such as low-frequency and structural variants (SVs) are poorly captured in association studies. To assess the impact of these variants on gene expression variation, we explored a half-diallel panel composed of 323 hybrids originated from pairwise crosses of 26 natural *Saccharomyces cerevisiae* isolates. Using short- and long-read sequencing strategies, we established an exhaustive catalog of single nucleotide polymorphisms (SNPs) and SVs for this panel. Combining this dataset with the transcriptomes of all hybrids, we comprehensively mapped SNPs and SVs associated with gene expression variation. While SVs impact gene expression variation, SNPs exhibit a higher effect size with an overrepresentation of low-frequency variants compared to common ones. These results reinforce the importance of dissecting the heritability of complex traits with a comprehensive catalog of genetic variants at the population level.

## Introduction

Gene expression variation among individuals corresponds to an essential step linking genetic variation and phenotypic diversity observed in natural populations (Hill *et al*, 2021; Albert & Kruglyak, 2015, 15; Rockman & Kruglyak, 2006). Dissecting the genetic basis of gene expression variation at the population level is therefore crucial to better understand the genotype-phenotype relationship. Genetic variants or loci associated with gene expression variation (*i.e*., expression Quantitative Trait Loci, eQTL) have been detected using different mapping strategies, such as linkage and genome-wide association studies (GWAS) (Mackay *et al*, 2009). Large-scale transcriptomic surveys in model and non-model systems have highlighted the preponderance of genetic variants impacting gene expression variation between individuals (Zhang *et al*, 2022; Albert *et al*, 2018; Kita *et al*, 2017; Schadt *et al*, 2003; Battle *et al*, 2014; West *et al*, 2007; Zhang *et al*, 2011; Ferraro *et al*, 2020; GTEx Consortium, 2017; Kawakatsu *et al*, 2016; Vu *et al*, 2015; Rockman *et al*, 2010; Caudal *et al*, 2023). In general, the transcript level of every gene appears to be influenced by one or more eQTL.

In humans, the genetic basis of gene expression variation was dissected via GWAS on a large dataset of 49 tissues obtained for up to 838 individuals (Ferraro *et al*, 2020). This study clearly highlighted tissue-specific gene expression patterns and detected a large number of eQTL across tissues. The detected variants are mostly local eQTL (*i.e*., located close to the genes they influence), while distant eQTL (*i.e*., located far from the genes they influence) remains difficult to identify in such context due to the large genomes, limited sample size and low statistical power. As most of gene expression variation does not arise from local eQTL, part of the variance is therefore still unexplored. More recently, a population-scale transcriptomic analysis was performed on more than 1,000 *Saccharomyces cerevisiae* yeast isolates leading to a deeper view of the genetic control of gene expression (Caudal *et al*, 2023). Overall, local eQTL were less frequent, representing 26% of the total set of eQTL detected, which is consistent with previous observation in a yeast cross (Albert *et al*, 2018). Nevertheless, the detected local and distant eQTL together only explained a small fraction of the gene expression heritability (Caudal *et al*, 2023), which is usually the case for all complex traits (Manolio *et al*, 2009; Hindorff *et al*, 2009). Several sources can potentially be considered as the origin of this missing heritability, *i.e*. the part of the phenotypic variance not unexplained by associated causal loci. These sources include the low power to detect small effect variants, to estimate non-additive effects (Cordell, 2009; Mackay, 2014; Zuk *et al*, 2012) but also the fact that rare and low-frequency variants (Manolio *et al*, 2009; Hindorff *et al*, 2009; Gibson, 2012; Pritchard, 2001; Walter *et al*, 2015) as well as structural variants (Peter *et al*, 2018) are not systematically taken into account in GWAS.

Unlike common variants, rare and low-frequency variants are not adequately captured or tested in standard GWAS analyses, as they are only present in less than 1% or 5% of the population, respectively. In humans, these variants have been shown to play a role and alter adult height for example (Marouli *et al*, 2017; Akiyama *et al*, 2019). Additionally, recent estimates of the impact of rare variants on the heritability for two human traits (height and body mass index) from whole-genome sequence data on 25,465 unrelated individuals of European ancestry confirmed that these variants are probably a major source of the missing heritability (Wainschtein *et al*, 2022). In the *S. cerevisiae* yeast model, the effect of rare and low-frequency variants on phenotypic variance has also been tested, since a bias towards this type of variants is observed, with more than 90% of the SNPs having a minor allele frequency (MAF) lower than 0.05 in a dataset of 1,011 yeast genomes (Peter *et al*, 2018). Based on this resource, two independent surveys have shown that these variants contribute disproportionately to growth variation in a large number of conditions observed across yeast natural isolates (Bloom *et al*, 2019, 19; Fournier *et al*, 2019). Although there is now strong evidence for the impact of such variants on organismal traits, their effect on molecular traits, such as gene expression variation, has been poorly studied at the population level.

Regarding the structural variants (SVs), they have long been recognized as an important source of genetic diversity and phenotypic variation in all organisms (Weischenfeldt *et al*, 2013). A few studies have provided catalogs of SVs in large populations of various organisms, including human, tomato, and yeast (Alonge *et al*, 2020; Li *et al*, 2023; O’Donnell *et al*, 2022; Liao *et al*, 2023), however, these catalogs are still non-exhaustive. Indeed, SVs are difficult to systematically and exhaustively detect for large natural populations and thus they are often excluded from complex trait association studies. In humans, certain SVs have been identified as involved in various diseases and are generally presumed to act through their effects on gene expression (Weischenfeldt *et al*, 2013). The impact of SVs on gene expression variation was estimated by exploring the effect of 61,668 SVs in 613 individuals on gene expression (Scott *et al*, 2021; Chiang *et al*, 2017). It was found that common SVs are causal for 2.66% of the eQTL and that SVs often affect multiple nearby genes (Scott *et al*, 2021). The impact of SVs on neighboring genes has also been recently studied in other organisms, such as tomato and yeast, and the results have shown variable effects on gene expression depending on the type of SVs (Alonge *et al*, 2020; O’Donnell *et al*, 2022). Although these studies represent the most comprehensive analysis of the impact of SVs on gene expression to date, they are still limited because either only part of the SVs are detected and not all SV-gene expression associations can be tested. Consequently, these studies do not provide a comprehensive view of the effect of SVs on the transcriptional landscape at the population level.

Understanding the effect of low-frequency variants as well as SVs on gene expression variation remains a challenge but it should provide deeper insights into the molecular basis of phenotypic diversity. Here, we took advantage of a diallel hybrid design in the *Saccharomyces cerevisiae* yeast model, which allows for exhaustive characterization of these two types of variants and their associations with gene expression variation in the population. Our findings show the need for a comprehensive catalog of genetic variants to identify the genetic basis of trait variation.

## Results

### Diallel design and transcript abundance variation

To better understand the sources of missing heritability of complex traits and more specifically of molecular traits such as transcript abundance, we sought to investigate the underlying genetic architecture of gene expression variation through the use of a diallel panel. In this context, a diallel cross panel was constructed by crossing a set of 26 genetically diverse *S. cerevisiae* natural isolates (Figure 1, Table S1). To capture a wide range of genetic diversity, different isolates from various ecological (*e.g*., beer brewery, fruit, clinical samples), and different geographical locations (*e.g*., Ireland, China, USA) were selected (Figure S1A-B). The pairwise nucleotide divergence between parental isolates ranges from 0.03% to 1.1%, with an average of 0.59% (Table S2). Parental isolates were crossed in pairs in all possible non-reciprocal combinations, a configuration often referred to as a non-reciprocal diallel cross. This diallel panel resulted in a total of 351 hybrids with 325 heterozygous hybrids coming from the cross of two different parents, and 26 homozygous hybrids resulting from the cross of genetically identical parental isolates.

**Figure 1.**
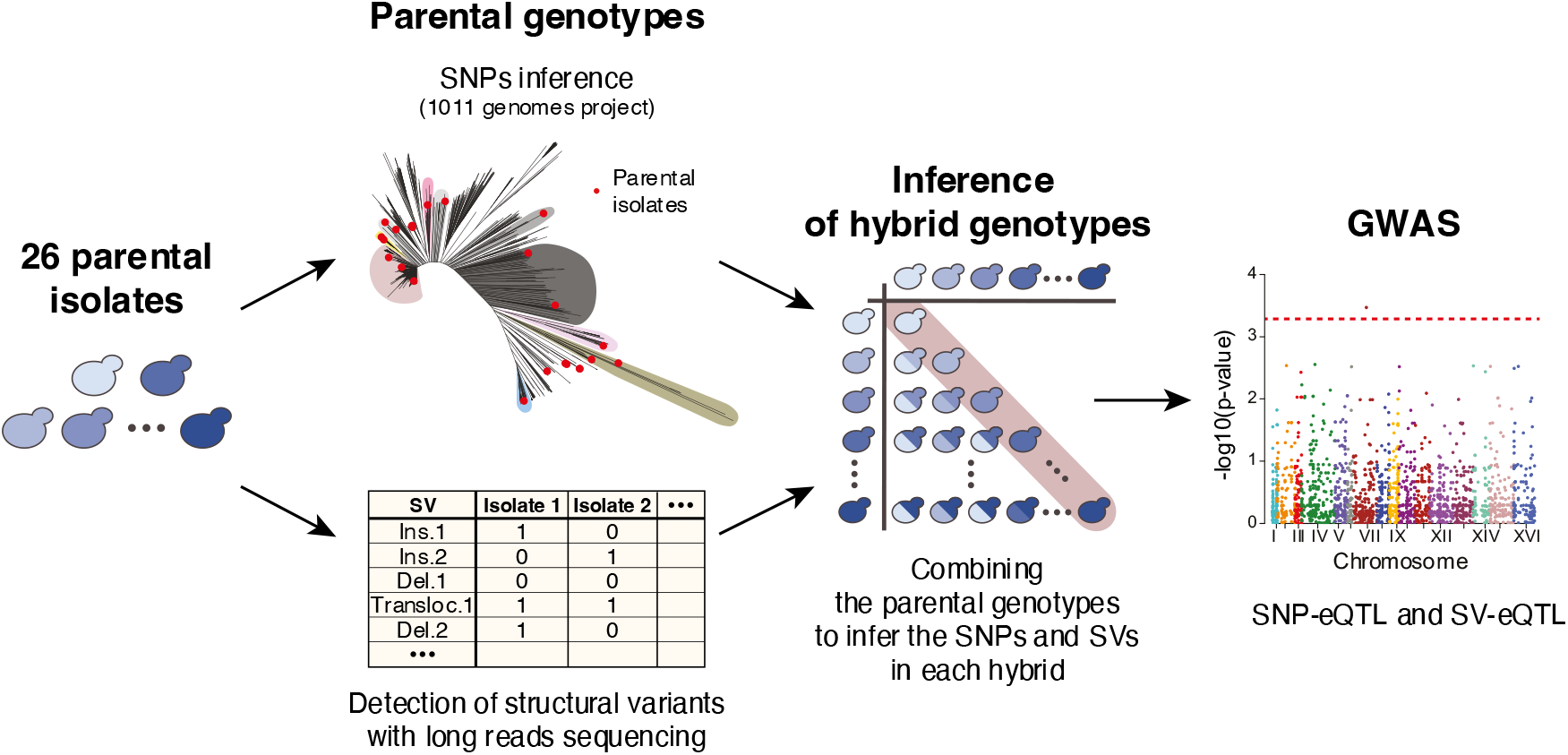
Workflow for genome-wide association using a diallel panel. A set of 26 natural isolates were selected as parents for a diallel panel. The SNPs in each isolate were inferred from the data of the 1,011 yeast genomes project (Peter *et al*, 2018) while their SVs were detected using long-reads Oxford Nanopore sequencing. The parental genotypes were combined pairwise to infer the SNPs and SVs in each of the hybrids produced by the diallel panel. Finally, the SNP and SV genotypes of the hybrids were used to perform GWAS on the transcript abundance of 6,186 genes.

To quantify gene expression variation in this population, RNA sequencing was performed on all the 351 hybrids (Tsouris *et al*, 2023). High-quality transcriptomes were obtained for 323 unique hybrids with over one million reads mapped to the reference genome. We obtained the expression levels (as transcript <u>p</u>er <u>m</u>illion or tpm) for 6,186 genes that are expressed in at least half of the samples (tpm > 0), including 5,740 core ORFs as well as 422 accessory ORFs that are variably present in the generated hybrids (Table S3). In total, we analyzed 2,002,550 transcriptomic measurements, providing a detailed analysis of the hybrid transcriptomes. The expression abundance (as the mean of log2(tpm+1)) and dispersion (as the mean absolute deviation of log2(tpm+1)) of each gene in the diallel population correlates with those determined in a large population of 969 *S. cerevisiae* natural isolates (*R*=0.79 and *R*=0.72, respectively) (Figure S1C-D) (Caudal *et al*, 2023). In addition, gene expression levels are also correlated between the homozygous hybrids generated in the current dataset and the original parental isolates that were characterized in the previously generated population-level dataset (Caudal *et al*, 2023) (*R*=0.73) (Figure S1E).

### Detection of the SNP-eQTL using the diallel panel

To determine the genotypes of all the hybrids, we retrieved the biallelic genetic variants of the 26 parental strains from the genomic sequences of the 1,011 yeast genomes project (Peter *et al*, 2018). As a diallel panel leads to a highly structured population, it may introduce some biases when performing genome-wide associations. Singletons, *i.e*. genetic variants that are only present in a parental genetic background, show a strong linkage throughout the genome. We therefore removed these variants from the genotype matrix, resulting in a SNP matrix of 31,818 variants with a minor allele frequency (MAF) higher than 5%. Using this genotype matrix, we quantified the genome-wide heritability (*h*^*2*^_*g*_) across the 6,186 gene expression traits (Figure S2A, Table S4). The median *h*^*2*^_*g*_ is 0.28, which is similar to heritability of gene expression estimated in linkage mapping and GWAS in yeast (median=0.26) as well as in other organisms such as in humans (mean=0.03-0.26) (Albert *et al*, 2018; Zhang *et al*, 2022; Huan *et al*, 2015; Ouwens *et al*, 2020).

To identify SNPs that influence gene expression variation (SNP-eQTL), we performed GWAS with the transcript abundance traits of 6,186 genes and using the SNP matrix as the genotype (Methods). We detected a total of 1,032 SNP-eQTL that influence the level of transcript abundance of 1,714 genes (Figure 2A, Table S5). Most SNP-eQTL impact only one or a few traits while few SNP-eQTL influence more than 20 phenotypes (Figure S2B). In fact, we detected a total of 4 eQTL hotspots located in the *BCY1, GAL2* and *MLP1* genes, while another one is located in the promoter of *GSH2* and are associated to the expression of 84, 66, 62, 57 genes, respectively (Figure 2B). The number of hotspots detected here is intermediate between the number identified by linkage mapping using single cross segregants and GWAS across a large set of natural isolates (Albert *et al*, 2018; Caudal *et al*, 2023). In addition, most traits are influenced by a single SNP-eQTL while very few are controlled by more than 5 (Figure S2C). Overall, a trait was associated to 1.77 eQTL while each eQTL influenced 2.94 different phenotypes on average.

**Figure 2.**
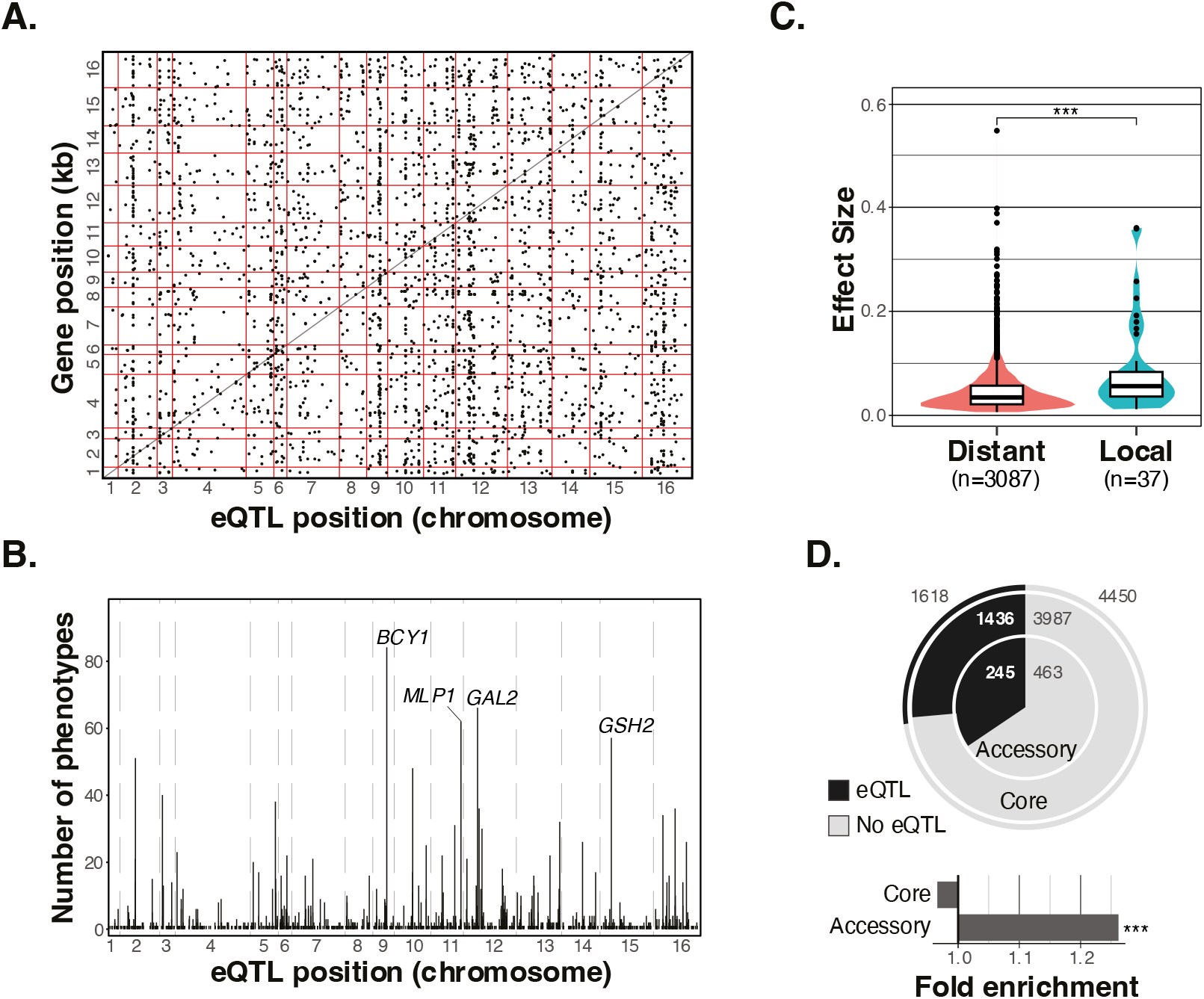
Genome-wide association results using the SNP matrix. (A) Positions of the SNP-eQTL and their associated genes along the genome. Each point represents an eQTL and the horizontal and vertical red lines mark the margins of the chromosomes. The diagonal black line indicates the positions where the eQTL position and gene position are the identical, therefore eQTLs on that line are very close to the gene they influence. (B) Positions of SNP-eQTL and the number of genes they are associated with. (C) Comparison of the effect sizes of distant and local-eQTL inferred from GWAS (two-sided Mann-Whitney-Wilcoxon test, p-value=3.9e^-4^). (D) The number and proportions of core (outer circle) and accessory genes (inner circle) associated with eQTLs. Fold enrichment of core and accessory genes associated to eQTLs. Significance was calculated using a two-sided Fisher test.

We then distinguished the SNP-eQTL according to their relative position with respect to the gene they impact, with local eQTL being close to the affected gene and distant eQTL located further away or on different chromosomes (see Methods). Consistent with previous observations, local SNP-eQTL display larger effect sizes compared to the distant ones (Wilcoxon test, p-value=3.9e-4) (Figure 2C) (Albert *et al*, 2018; Caudal *et al*, 2023).

Gene expression regulation of accessory genes has been shown to differ from that of the core genome (Caudal *et al*, 2023). We therefore compared the likelihood of accessory and core genes to be associated to SNP-eQTL (Figure 2D). Accessory genes are more likely to be associated with SNP-eQTL (Fisher test, p-value=3.7e-3). In total, 34.6% of the accessory genes (245 out of 708 accessory genes) considered in GWAS are associated with an eQTL, compared to a 26.7% of all genes (1,618 out of 6,068 genes), with a 1.26-fold enrichment. In addition, SNP-eQTL associated with accessory genes have a larger effect size on average than those associated with core genes (Wilcoxon test, p-value=3.7e-6). Specifically, the effect size of distant SNP-eQTL controlling accessory genes is larger than those controlling core genes (Wilcoxon test, p-value=3.96e-6). However, no significant difference was observed between the local SNP-eQTL associated with the expression of accessory and core genes (Wilcoxon test, p-values=0.6) (Figure S2D).

### Impact of the low-frequency SNPs on gene expression variation

Next, we sought to explore the contribution of rare and low-frequency genetic variants (*i.e*., with a MAF<0.01 and <0.05, respectively) to the observed gene expression variation. In fact, genetic variants considered by GWAS must have a relatively high frequency in the population to be detectable, usually over 0.05 for relatively small datasets (Visscher *et al*, 2017). Rare and low-frequency variants are therefore excluded from association studies. However, our diallel panel provides a powerful strategy to assess the impact of low-frequency variants that are initially present in the population, *i.e*. in the 1,011 *S. cerevisiae* collection here (Peter *et al*, 2018). In fact, each parental genotype is present several times, increasing the frequency of these variants in this new population and allowing their detection in GWAS. While the parental genomes carry 1,205 low-frequency variants (81 of which are rare variants) as defined in the population of 1,011 isolates, none of the genetic variants has a MAF lower than 0.058 in our diallel panel (Datafile 1).

Interestingly, the MAF of the genetic variants identified as SNP-eQTL is much lower than that of the entire set of SNPs considered for GWAS, suggesting that low-frequency variants have a large impact on transcript abundance variation (Figure 3A). In fact, 10.1% (a total of 104 SNPs) of the significant SNP-eQTL have a MAF below 0.05, whereas only 3.6% (a total of 1,141) of SNP matrix used for GWAS have a MAF below 0.05 in the natural population (Figure 3B). This nearly 3-fold enrichment in low-frequency SNPs in the set of associated variants is consistent with previous studies focusing on the influence of low-frequency variants on yeast growth traits (Bloom *et al*, 2019; Fournier *et al*, 2019) (Figure 3C). Using a diallel panel, we previously have found that 16.8% of growth SNP-QTL were low-frequency variants in the collection of 1,011 *S. cerevisiae* isolates (Fournier *et al*, 2019). Finally, we measured the respective effect size of each SNP-eQTL and we found that even if enrichment is observed, low-frequency variants have a lower effect size compared to the common variants (Figure 3D).

**Figure 3.**
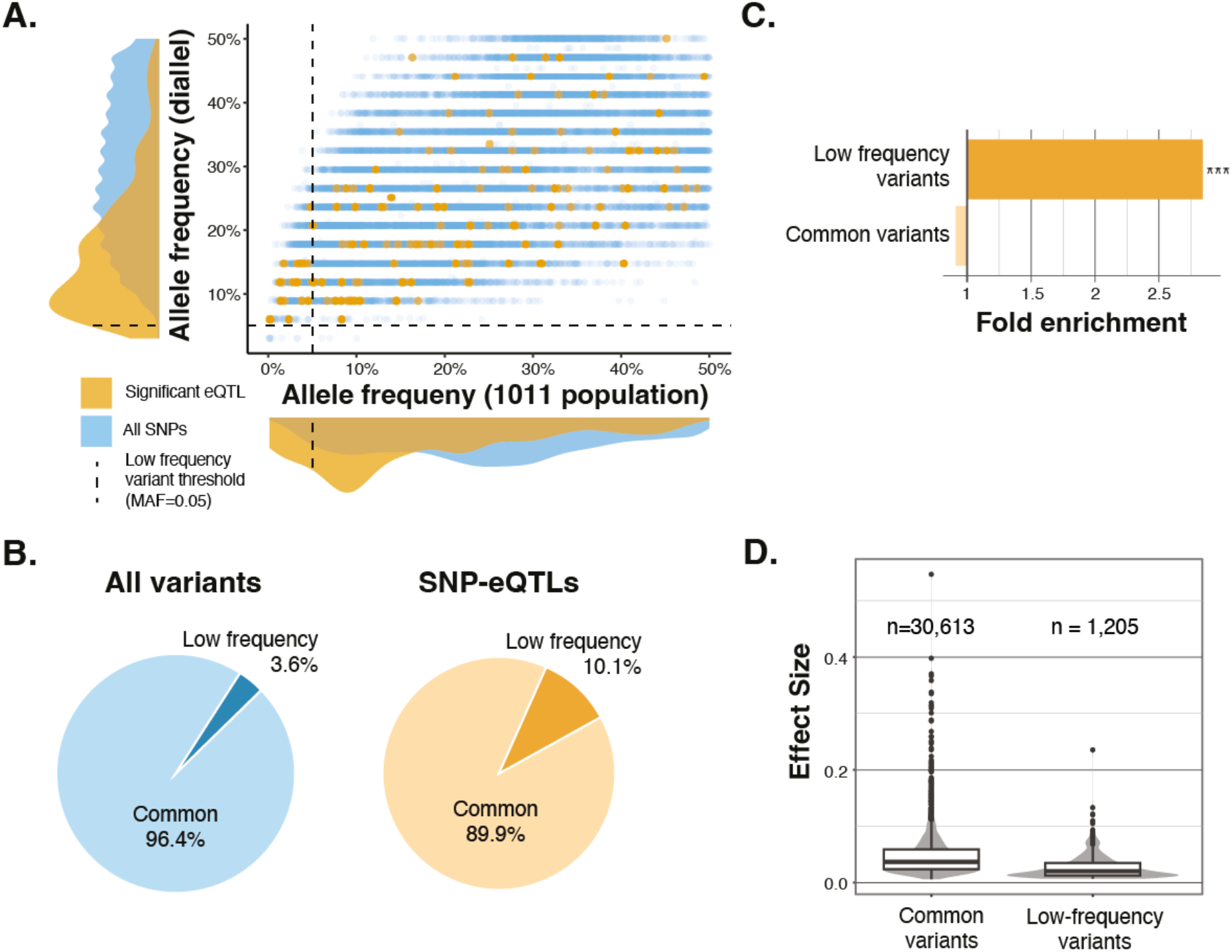
Low frequency variants and gene expression variation. (A) Comparison of the MAF of the SNPs used in GWAS in the diallel population and the natural population (1011 isolates). Orange points represent significant eQTL SNPs whereas blue points represent the remaining SNPs. In the distributions along the axes, the orange distribution covers the significant SNPs, and the blue distribution covers all the SNPs used in GWAS. Vertical and horizontal dashed lines show the threshold of MAF (5%) used to define low-frequency variants. (B) Comparison of the fraction of low frequency variants in all the SNPs used in GWAS (blue) and the significant eQTLs (orange). (C) Fold enrichment of low frequency and common variants to be detected as SNP-eQTL (two-sided Fisher test, p-value<2.2e^-16^). (D) Comparison of the effect size of common and low-frequency SNP-eQTL

### Characterization of the structural variants in the diallel panel

We then focused on the impact of SVs on gene expression variation using our diallel panel. Accurate and exhaustive identification of SVs in a large population can be very laborious and, in some cases, practically impossible. These variants are therefore only very rarely taken into account in association studies. However, the structure of a yeast diallel panel makes this task much easier because the SVs present in the hybrids correspond to the combination of the SVs present in their parents. Long-read genome sequencing and SV detection thus provide an exhaustive list of the SVs present in each of the 323 hybrids.

We therefore sequenced the genomes of 24 out of 26 parental isolates using a long-read Oxford Nanopore sequencing strategy, as two of the parental strains, S288C and ∑1278b, were already completely sequenced and assembled (Goffeau *et al*, 1996; Dowell *et al*, 2010). On average, we obtained 108,351 reads with a length of 12.8 kb for each parental genome (Table S6). The mean coverage was 66X on average, ranging from 19.7X to 150X for the ABA and ACS isolates, respectively. We then used these long reads to generate *de novo* assemblies for the entire set of strains (see Methods). Overall, we obtained contiguous assemblies with a N50 of 835 kb and an average genome size of 12.2 Mb (Table S7). In most cases, a single contig covered more than 90% of a chromosome, while in other cases, only a few contigs were needed. (Figure S3A). The assembly of each parent was then compared to the genome of the reference strain (S288C) using *MUM&co* to detect all the SVs present in the parental genomes (O’Donnell & Fischer, 2020). As the SVs are detected independently in each parent, we also had to identify the presence of the same event in the several genetic backgrounds in order to merge them into the same occurrence. For this purpose, we used *Jasmine* (Kirsche *et al*, 2023), a tool that compares the genomic position as well as the sequences of SVs, and we found a total of 1,953 SVs in our set of parental genomes (Table S8).

The SVs were classified into 6 types, namely insertions (n=1,032 variants), deletions (n=543), duplications (n=250), contractions (n=65), translocation (n=33) and inversions (n=30) (Figure 4A). The number of SVs in each parent correlates with the nucleotide divergence between that parent and the reference genome (*R*=0.71, p-value= 7.30e^-05^) (Figure S3B). While ∑1278b has the lowest nucleotide diversity compared to S288C and the lowest number of SVs (n=153), the BAM isolate has the highest number of SVs (n=357) and corresponds to the most divergent strain with respect to S288C (Figure 4B). On average, we detected 241 SVs per genome. The distribution of SV types in each parent is similar to their overall abundance. Most SVs are insertions and deletions, while the remaining SVs types only represent a small fraction. As with SNPs, SVs exhibit a bias towards low-frequency alleles (Figure S3C). A total of 60.1% of SVs (n=1,190) are low-frequency variants, with only 763 SVs have a MAF greater than 0.05 (Figure S3C).

**Figure 4.**
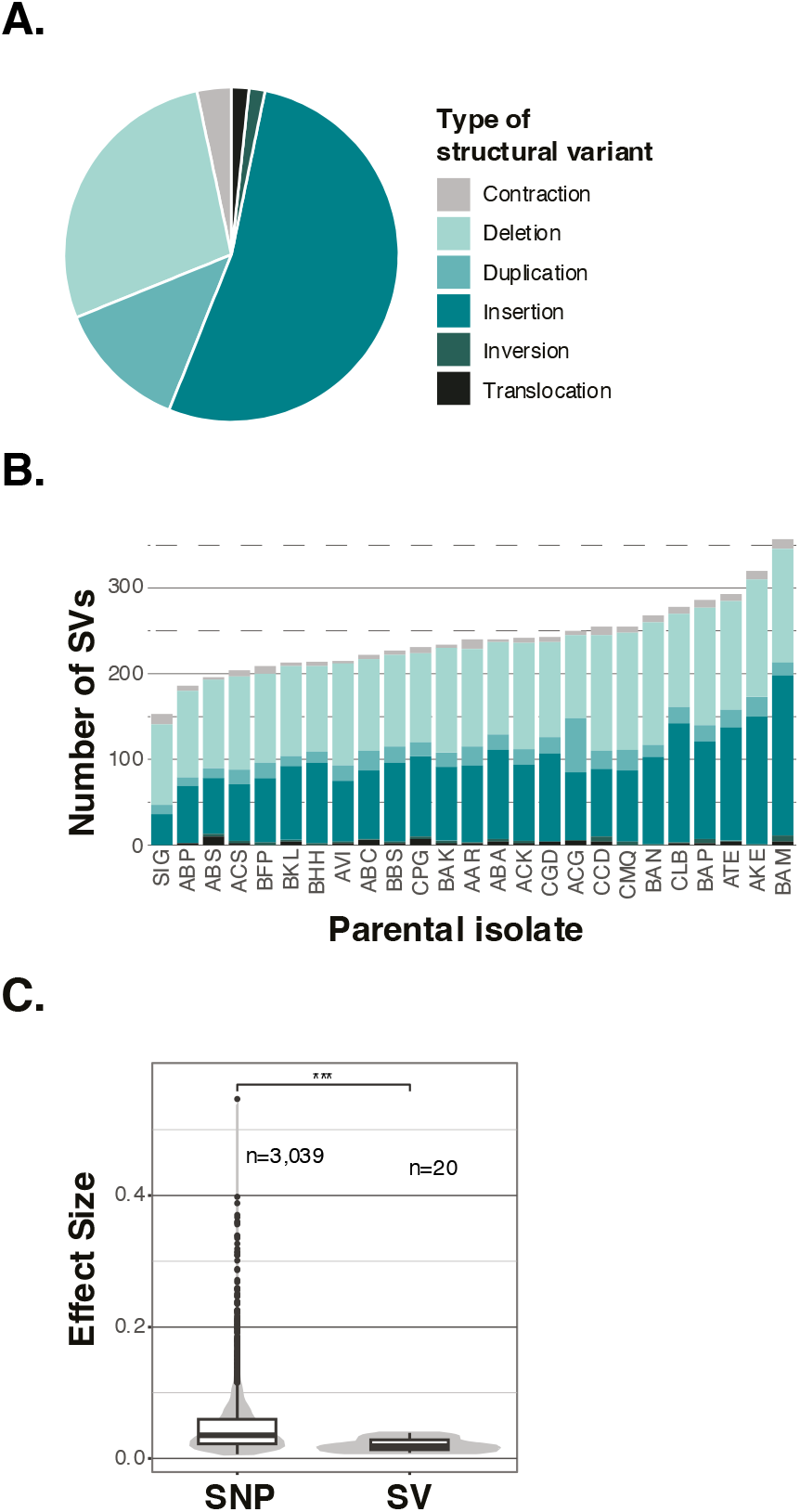
Structural variants detected in the parental genomes and effect size comparison. (A) Fractions of the different types of structural variants identified in the parental strains. (B) Number of the SVs detected in each parental strain. Each parent is pictured in each stacked bar and colors of the bars represent the different types of SV as above. (C) Comparison of the effect sizes of SNP-eQTL and SV-eQTL identified through GWAS (two-sided Mann-Whitney-Wilcoxon test, p-value=6.48e^-12^).

Finally, we also identified SVs that are related to transposable elements, a major source of structural variation (see Methods). Overall, 869 SVs are *Ty*-related (44.5%), while 1,084 (55.5%) are not (Figure S3D). *Ty*-related SVs can be separated into two groups according to their length. The first group of SVs are about 340 bp in length and contains the *Ty* Long Terminal Repeats (LTR), while the second group of SVs are about 6 kb in length and contains the entire or partial *Ty* elements (Figure S3D). A total of 64.2% (n=558) of *Ty* related SVs only contain LTR sequences, while the remaining 35.8% (n=311) contain partial or complete *Ty* elements.

### Impact of SVs on gene expression phenotypes

Before performing genome-wide associations, we inferred the genotypes of the 323 hybrids by combining the SVs of their parents as it was done previously for the SNPs. We excluded the singletons, *i.e*. all variants present in only one of the 26 parents, because they would cause many false positive associations due to severe linkage throughout the genome. As a result, we obtained a genotype matrix of 763 SVs across the 323 hybrids. The *h*^*2*^_*g*_ of transcript abundance traits using the SV genotype matrix was 0.021, an order of magnitude lower than the *h*_*2*_*g* obtained with the SNP matrix (*h*^*2*^_*g*_=0.28) (Figure S4A).

To identify SVs that influence gene expression variation (SV-eQTL), we performed GWAS with the transcript abundance traits of 6,186 genes and using the SV matrix as the genotype (see Methods). We only detected 19 significant SV-eQTL that influence the expression of 13 genes (Table S9). The transcript abundance of 10 genes was influenced by one SV-eQTL, while the expression of two genes was influenced by multiple SV-eQTL (Figure S4B). All SV-eQTL are associated to a single gene except for one that was associated to the transcript abundance of two genes (Figure S4C). Most SV-eQTL are insertions and deletions (n=13 and n=4, respectively), while one SV-eQTL is a duplication located on chromosome 13 and another one a contraction on chromosome 15. More than half of the SV-eQTL are *Ty*-related (10 out of the 19 SV-eQTL) but this is not significant compared to the overall abundance of *Ty*-related SVs (fisher test, p-value=1). Finally, since we assessed the relationship of SNPs and SVs to the same phenotypes across the same population, it was possible to compare the effect of SNPs and SVs on gene expression. In fact, we found that the effect sizes of SV-eQTL are overall smaller than those of SNP-eQTL (Figure 4C).

## Discussion

Understanding the genetic basis of gene expression variation is essential to have a better insight into the genotype-phenotype relationship. While large-scale studies detected thousands of genetic variants involved in gene expression variation via genome-wide associations, these variants explain only a small fraction of gene expression variation (Ferraro *et al*, 2020; GTEx Consortium, 2017). In this context, we sought to investigate the impact of potential sources of this so-called missing heritability. Using a diallel design, we were able to assess the influence of low-frequency as well as structural variants on the variation in transcript abundance. On the one hand, this strategy makes it possible to artificially increase the frequency of genetic variants in a diallel panel and thus to take low-frequency variants into account in association studies. On the other hand, it is possible to easily establish the exhaustive list of SVs present in the hybrids from the parental genotypes. Based on a generated yeast diallel panel composed of 323 hybrids and associated with the corresponding transcriptomes, we carried out a GWAS for each gene expression trait and identified 1,032 SNP-eQTL and 19 SV-eQTL in total.

Regarding the low-frequency and rare variants, we found a strong enrichment of these variants in the set of significantly associated SNPs, *i.e*., SNP-eQTL. While only 3.6% of the SNPs considered in the association tests have a low frequency in the natural yeast population, we found that 10.1% of all SNPs-eQTL are low-frequency variants, showing a 3-fold enrichment. These results clearly show the same trend and enrichment already observed for yeast growth phenotypes (Bloom *et al*, 2019; Fournier *et al*, 2019). This observation highlights that low-frequency variants strongly contribute to gene expression variation in a natural population. This is very important as much of the detected genetic polymorphisms in a population, such as the 1,011 yeast genomes dataset, are low-frequency variants with almost 92.7% of the polymorphic sites associated with a MAF lower than 0.05 (Peter *et al*, 2018). Nevertheless, it is also worth mentioning that alongside this enrichment, we also found that overall, the effect sizes of these variants are smaller than those of the common variants.

Our diallel panel, however, showed some limitations in exploring the impact of SVs on gene expression variation. In fact, we could only detect a total of 19 SV-eQTL probably because very few SVs were taken into account compared to SNPs in the genotype matrices (763 SVs *versus* 31,818 SNPs). By taking into account the size of the genotypic matrices, we can observe that the same fraction of SNP-eQTL and SV-QTL was globally detected, corresponding respectively to 3.3% and 2.6% of associated genetic variants, respectively. It is clear that SVs are widely present in natural populations, as in *S. cerevisiae* for example. A recent study of 142 telomere-to-telomere genomes from natural isolates characterized approximately 4,800 unique SVs and identified 97 SVs that influence the expression of neighboring genes (O’Donnell *et al*, 2022). The impact of SVs on gene expression has also been shown in humans where SVs are responsible for 2.66% of eQTL and they often affect multiple nearby genes (Scott *et al*, 2021). The impact of SVs is not limited to gene expression traits, a survey of over 100 tomato genomes has indeed identified hundreds of SV-QTLs impacting several volatile flavor and fruit metabolites (Alonge *et al*, 2020; Li *et al*, 2023).

Our diallel design also comes with other limitations. First, the diallel hybrid population is very structured, and in order to avoid biases we had to eliminate many variants that were in linkage, therefore limiting the number of variants used in GWAS. This limit could possibly be circumvented by significantly increasing the size of the panel. Second, the hybrids were not sequenced directly but their genotypes were inferred from their parents. Therefore, any potential genomic modifications that have arisen since the formation of the hybrids were generated (such as any mutation, loss of heterozygosity or other structural events) are not considered in our design. Finally, as already mentioned, the number of SVs used for the association studies was probably insufficient to have a global view of the impact of SVs on gene expression.

A population-scale comprehensive survey would now be essential to have an accurate and exhaustive view of the impact of SVs on gene expression variation. The collection of 1,011 natural yeast isolates could be an excellent resource for such exploration. Long-read sequencing of this large collection would lead to a species-wide view of the structural variants as well as a comprehensive pangenome graph (Liao *et al*, 2023; Hickey *et al*, 2023). The exhaustive catalog of SVs generated could then be used to perform GWAS on a large number of traits. Many traits have already been measured for this large collection of *S. cerevisiae* isolates, including growth phenotypes, as well as molecular traits like transcript and protein abundance (Caudal *et al*, 2023; Peter *et al*, 2018; Muenzner *et al*, 2022; Teyssonniere *et al*, 2023). A better view of the impact of SVs on traits would definitely improve our understanding of the genotype-phenotype relationship by revealing their real role in missing heritability.

## Material and methods

### Diallel panel generation and cell harvesting

The hybrid diallel panel was produced from a series of pairwise crosses between 26 haploid parental strains. Two stable haploid lines were generated for each parental strain, the *MATa* lines carry a *KanMX* cassette and the *MATalpha* lines carry a *NatMX* cassette, each cassette replacing the *HO* locus (Fournier *et al*, 2019). For each cross cells of opposite mating type were patched together on solid YPD media (1% yeast extract, 2% peptone and 2% glucose) and incubating them at 30°C overnight. To select for hybrid cells by selecting the combination of both cassettes the cell patches were then transferred to YPD media containing G418 (200 mg/ml) and nourseothricin (100 mg/ml) and incubated at 30°C overnight. All procedures were done using the replicating robot ROTOR (Singer Instruments). In total, we obtained 351 genetically unique hybrids.

Hybrids were then incubated in liquid synthetic complete (SC) media with 2% glucose in 96-deep-well plates for cell harvesting. The optical density of each hybrid’s culture was measured systematically using a 96-well microplate reader (Tecan Infinite F200 Pro). During log-phase growth (OD_600nm_ ∼0.3), cells were harvested by filtration in 96-well filter plate (Norgen, #40008) where the media was eliminated by applying vacuum (VWR, #16003-836). The filter plates were immediately flash frozen in liquid nitrogen and stored at -80°C.

### cDNA library preparation

We used the Dynabeads® mRNA Direct Kit (ThermoFisher #61012) to extract the mRNA of each hybrid. The cells were lysed using glass beads and were then incubated at 65°C for 2 minutes. mRNA was selected with two rounds of hybridization of their polyA tails to magnetic beads coupled to oligo(dT) residues.

cDNA sequencing libraries were prepared with the NEBNext® Ultra™ II Directional RNA Library Prep Kit (NEB, #E7765L) and using the manufacturer’s protocol. The concentration of cDNA in each library was quantified using the Qubit ™ dsDNA HS Assay Kit (Invitrogen ™) in a 96-well plate using a microplate reader (Tecan Infinite F200 Pro) with an excitation frequency of 485 nm and emission of 528 nm. Fragment size was assessed with Bioanalyzer 2100 (Agilent™) using the High sensitivity DNA kit (#5067-4626). We generated sequencing pools containing equimolar fragments from each sample. Lastly, the pools were sequenced for 75 bp single-end with Nextseq 550 (Illumina™) sequencer at the EMBL Genomics Core Facility.

### mRNA abundance quantification

For each sample the raw sequencing reads were mapped to a custom reference genome using STAR (Dobin *et al*, 2013) with the following parameters:

~~~
--outSAMtype BAM SortedByCoordinate \
--outFilterType BySJout \
--outFilterMultimapNmax 20 \
--outFilterMismatchNmax 4 \
--alignIntronMin 20 \
--alignIntronMax 2000 \
--alignSJoverhangMin 8 \
--alignSJDBoverhangMin 1
~~~

The custom reference genome combines the 16 chromosomes of the R64_nucl reference genome with the accessory ORFs present in the parental strains (n=665) according to the data from the 1011 yeast genomes project(Peter *et al*, 2018), each accessory ORF is considered as a separate contig. In total, 323 hybrids had enough reads and were used in the subsequent analyses. The reads aligning to each gene of the reference (n=6,285) and accessory (n=665) genomes were counted using the featureCounts function of the R package subread (Liao *et al*, 2014) with the parameter *countMultiMappingReads=F* in order to not take multi-mapped reads into account. For a given hybrid, if accessory genes that have orthologs in the reference genome were annotated as absent, we merged their reads counts to those of their reference genome counterparts. mRNA abundance was then normalized by calculating the transcripts per million value (tpm) of each gene. This gave us a list of tpm values for 6917 genes. We filtered out genes that have a zero tpm value in more than half of all samples, leading to a final dataset of 6186 genes (Tsouris *et al*, 2023).

### Long-reads sequencing of the parental genomes

To carry out long-reads sequencing of the parental isolates with Oxford Nanopores Technologies (ONT) sequencing, we first incubated each parental isolate in 40 ml of YPD (1% yeast extract, 2% peptone and 2% glucose) at 30°C for 48 hours. The cells were then harvested by centrifugation at 7,000 g for 3 minutes and then resuspended water before a second round of centrifugation. The cell pellet was resuspended in 4mL of sorbitol 1M containing 250 μl of zymolyase (1000 U/ml) and incubated at 30°C for 2 hours with agitation. Protoplasts were recovered by centrifugating for 5 minutes at 2,000 g at 4°C and discarding the supernatant. They were then resuspended in 4mL of lysis buffer, a solution of Tris-HCl 125mM, EDTA 62.5 mM, NaCl 0.625 M, 0.05 g PVP40, 1.25% SDS and 10 mg/ml RNaseA. The cell lysis suspension was incubated at 50°C for 3 hours and was then chilled on ice for 2 minutes. We then added 5 ml TE and 3 ml AcK 5 M to the suspension and centrifugated twice at 2,000 g for 15 minutes at 4°C. After each centrifugation the supernatant was transferred to a new Falcon tube where we then added 12 ml of isopropanol to precipitate the genomic DNA. After a 5-minute centrifugation at 500 g the DNA pellet was washed with ice cold 70% ethanol and incubated for 5 minutes on ice. To precipitate the DNA pellet, we carried out a centrifugation of 5 minutes at 500g at 4°C and then removed all ethanol from the tube. Finally, we added 250 μl of TE without disturbing the pellet and incubated overnight at room temperature before recovering the resuspended DNA without recovering any of the remaining pellet.

Some parental isolates were sequenced in-house using MinION R9.4.1 flowcells (10 isolates) whereas 14 isolates where sequenced by the Genoscope using PromethION R9.4.1 flowcells. All libraries were prepared with the Ligation Sequencing Kit SQK-LSK109. In order to sequence multiple isolates in the same run, the isolates sequenced with the MinION and PromethION were barcoded with the EXP-NBD104 and NBD114.96 Native Barcoding Kits respectively. Basecalling was carried out with guppy 5.0.16 (MinION data) and 5.0.11 (PromethION data).

### Genome assembly and structural variant detection

Raw fastq files were processed using Porechop (v.0.2.4; github.com/rrwick/Porechop) in order to remove adapters and barcodes. We further treated these reads using Filtlong (v.0.2.1; github.com/rrwick/Filtlong) to remove reads shorter than 1000 nucleotides. For each strain, reads were then assembled using both Canu (v. 2.2) (Koren *et al*, 2017) and SMARTdenovo (Liu *et al*, 2021). Canu assemblies were generated using the options *-nanopore-raw* and *genomeSize=12m* while we used the default options for SMARTdenovo. We used the same fastq reads to polish both assemblies per strain using Medaka (v. 1.7.2; github.com/nanoporetech/medaka).

We then set out to detect SVs using MUM&Co (v 3.8) (O’Donnell & Fischer, 2020) separately on each genome assembly, using the option *-g 12000000*. Unique identifiers were also added to each SV using a combination of bcftools (Danecek *et al*, 2021) and the R package vcfR (v1.12) (Knaus & Grünwald, 2017). We then used Jasmine (Alonge *et al*, 2020; Kirsche *et al*, 2023)(to collapse SV calls during two rounds. First, for each strain, SV calls from both Canu and SMARTdenovo assemblies were collapsed into 24 strains specific VCF files. During the second round of merging, the 24 VCF files and a 25^th^ containing direct SV calls from the publicly available ∑1278b genome (Dowell *et al*, 2010) were collapsed into one final VCF file for downstream analyses. Then, since the 26^th^ parent is a haploid version of the reference strain we assigned all the SVs as absent in that parent. Finally, SVs linked to transposable elements were detected using BLAST (Johnson *et al*, 2008)and a database containing LTR and Ty sequences (Bleykasten-Grosshans *et al*, 2021). Any SV covered by Ty-related elements on more than 50% of its length was flagged as Ty-related.

### Generation of the genotype matrix

We recovered the genotypes for all the hybrids in our diallel panel from Fournier *et al*., 2019 where the parental genotype had been combined to infer the genotypes of the hybrids (Fournier *et al*, 2019). We retained biallelic variants using vcftools with the option *--min-alleles 2* and excluded singletons with the vcftools *--singleton* and *--exclude* commands, resulting in a final matrix of 31,818 SNPs. The genotype matrix was then recoded to a bed file with the ‘recode12’ function of PLINK (Chang *et al*, 2015). The SVs were detected using MUM&co were subjected to the same filtering and recoding process, in order to be used for GWA analyses. The gene expression phenotypes, measured as transcripts per million, of 6186 genes were recovered from Tsouris *et al*., 2023 (Tsouris *et al*, 2023). The z-scores of each phenotype were then calculated and later used in GWAS.

### Genome-wide heritability estimation

The genotype matrices used to estimate the genome-wide heritability were generated from the genotype data of the 1011 yeast genomes project (for the SNP matrix) and from the SVs identified using long-reads sequencing (for the SV matrix). To decrease linkage between the individuals we removed the variants present in only one parent. Furthermore, the variants with strong linkage disequilibrium (r^2^ > 0.8) were removed for the calculation of the genome-wide heritability estimation. We calculated the weights of each variant using ldak with the –cut-weights and –calc-weights-all arguments and the default parameters(Zhang *et al*, 2021). All variants with non-zero weights were to generate a filtered vcf matrix of 5,493 SNPs that was then recoded with the plink -make-bed command. The filtered and recoded matrix was used to calculate the kinship between the individuals using the popkin function of the R package popkin with the default parameters. To estimate genome-wide heritability (*h*^*2*^_*g*_), from the kinship matrix mentioned above we used the hglm R package (Rönnegård *et al*, 2010) using the default parameters (Tsouris *et al*, 2023).

### Genome-wide association

Genome-wide association analyses on transcript abundance z-scores were performed with the single_snp function of the FaST-LMM python package (Widmer *et al*, 2014). We calculated condition-specific p-values by permuting the phenotypic values of the individuals 100 times and setting average 5% quantile (5% lowest p-values) as the threshold. This method doesn’t provide us with a false discovery rate (FDR). Independent GWA analyses were carried out for SNPs and SVs but the p-value thresholds for the GWA on SNPs and on SVs were very different due to the important difference between the number of SNP and SV used. To correct for this discrepancy, we adjusted the condition-specific p-values of each GWA by normalizing to the total number of variants (SNPs and SVs).

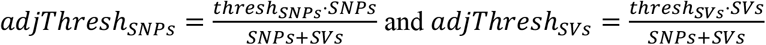

The GWA eQTL displayed some genetic linkage associated patterns/signatures driving groups of linked variants to pass the significance threshold. We calculated genetic linkage between variants using the plink --r command (Chang *et al*, 2015) and for each linkage group with an *R* value over 0.8 we only retained the variant with the lowest p-value. This variant filtering steps were independently run for the GWASs using the SNP and SV.

To distinguish between local and distant eQTLs we used a threshold of 25 kb, where the variants withing 25 kb of the start site of a gene were considered as local whereas those further away were considered as distant. We tested the differences in effect sizes or absolute variant weight between groups of eQTLs with Wilcoxon-Mann-Whitney tests using *wilcox.test* R function with the default parameters.

### Analysis of low frequency variants

The MAF of the SNPs in the hybrids and the natural population of 1,011 isolates was calculated with the vcfR using the *maf* command (Knaus & Grünwald, 2017) using the genotype matrix that was used for GWAS and the genotype matrix of the 1,011 yeast genomes project respectively. The fold enrichment values of eQTLs from low frequency and common variants were carried out manually in R and tested using the *fisher.test* command.

## Data availability

All Oxford Nanopore sequencing reads are available in the European Nucleotide Archive (ENA) under the accession number PRJEB64478.

The 1002 Yeast Genome website – http://1002genomes.u-strasbg.fr/files/diallel_RNAseq provides access to:

- Datafile 1: MAF of the SNPs across the diallel population and the population of 969 natural isolates.

## Acknowledgments

This work was supported by a National Institutes of Health (NIH) grant R01 (GM147040-01), a European Research Council (ERC) Consolidator grant (772505) to J.S and a French National Research Agency (ANR) young investigator grant (ANR-22-CE12-0023-01) to J. H. It is also part of Interdisciplinary Thematic Institutes (ITI) Integrative Molecular and Cellular Biology (IMCBio), as part of the ITI 2021-to-2028 program of the University of Strasbourg, CNRS, and Inserm, supported by IdEx Unistra (ANR-10-IDEX-0002). J.S. is a Fellow of the University of Strasbourg Institute for Advanced Study (USIAS) and a member of the Institut Universitaire de France.

