## Supplemental Material for "Diallel panel reveals a significant impact of low-frequency genetic variants on gene expression variation in yeast"

for

**This document includes:**

Supplementary figures S1-S4

Supplementary figure legends

Description for supplementary tables S1-S9x

Online Datafile descriptions

**Figure S1**

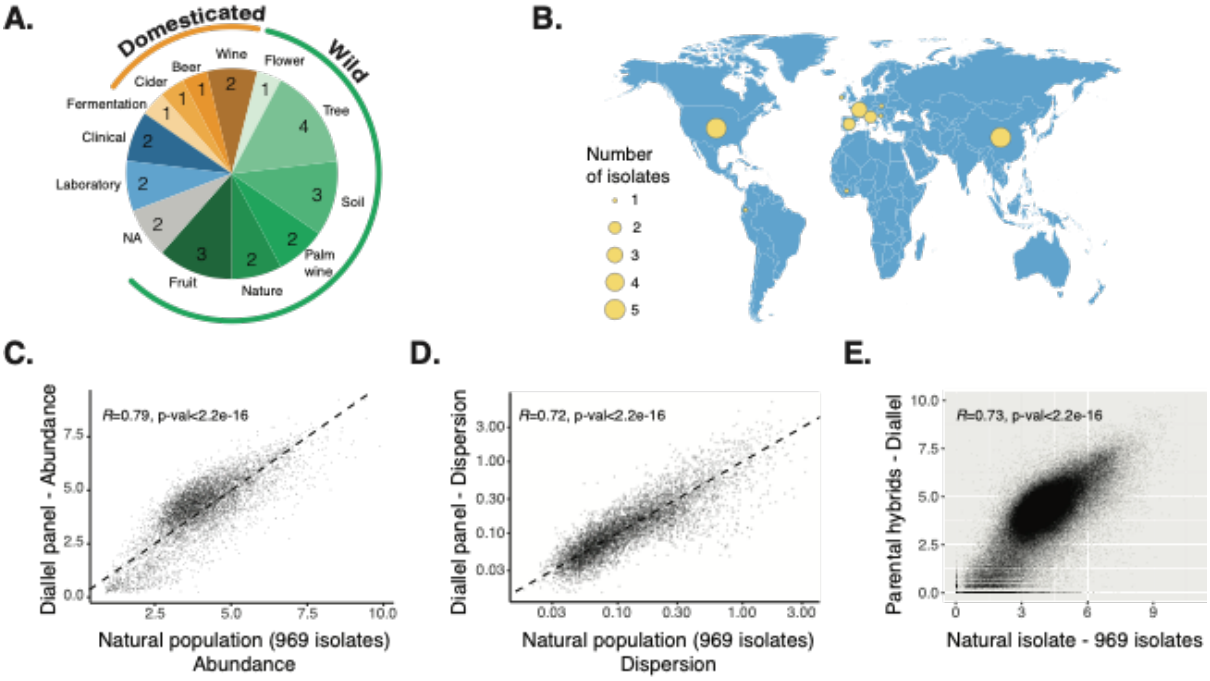


**Figure S1. Selection of the parental isolates and correlation between the transcript abundance in the diallel and the natural population**

(A, B) Ecological and geographical origin of the parental isolates that were used to generate the diallel crossing panel. Correlation of the average transcript abundance (C) and dispersion (D) for each gene between the diallel population and the population of 969 natural isolates. (E) Correlation of the transcript abundance of each homozygous hybrid and the respective homozygous diploid natural isolate.

**Figure S2**

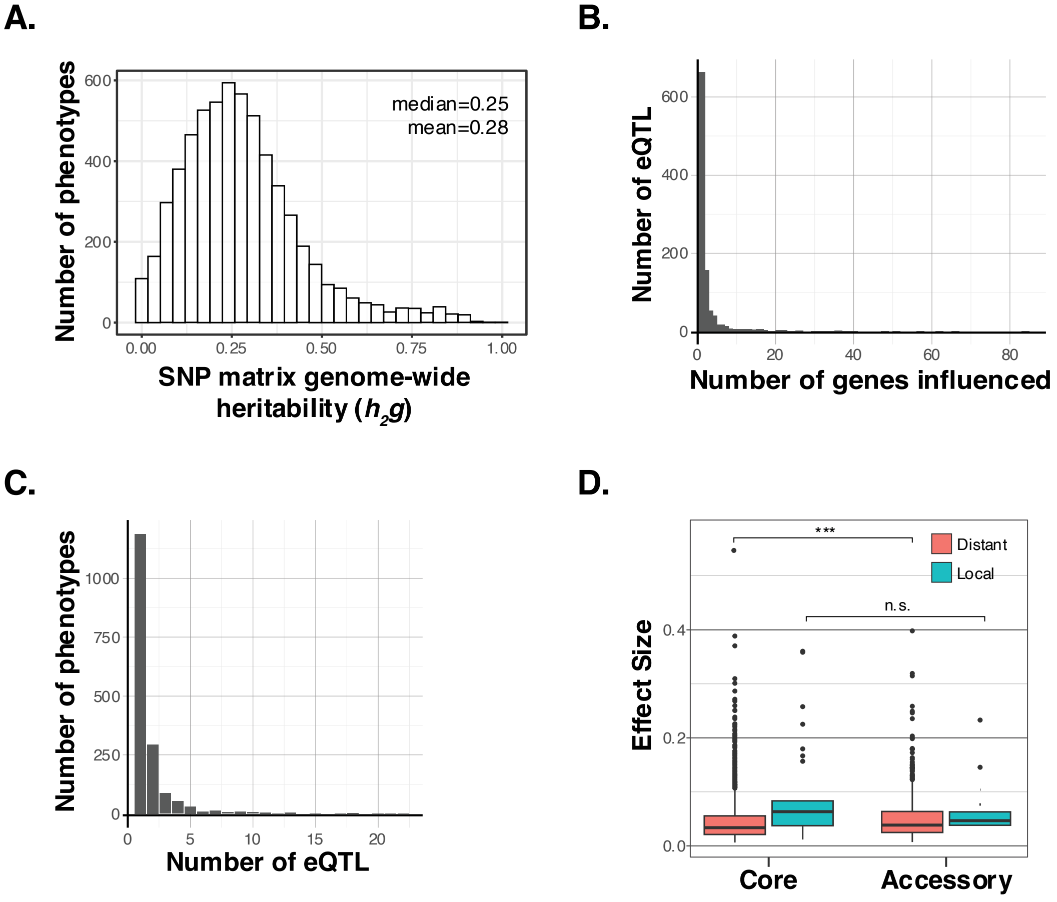


**Figure S2. Genome-wide heritability and GWAS results using the SNP**

(A) Distribution of genome-wide heritability ($h^2g$) of all transcript abundance phenotypes calculated based on the kinship matrix of the hybrids. (B) Distribution of the number of genes influenced by each SNP-eQTL. (C) Distribution of the number of eQTLs associated to each gene. (D) Effect size of eQTLs based on their position, distant or local in red and cyan, and the type of gene that is influenced (core or accessory gene). Significant difference is observed only between the distant eQTLs impacting core and accessory genes (two-sided Mann-Whitney-Wilcoxon test, p-value=$2.4e^{-4}$).

### Figure S3

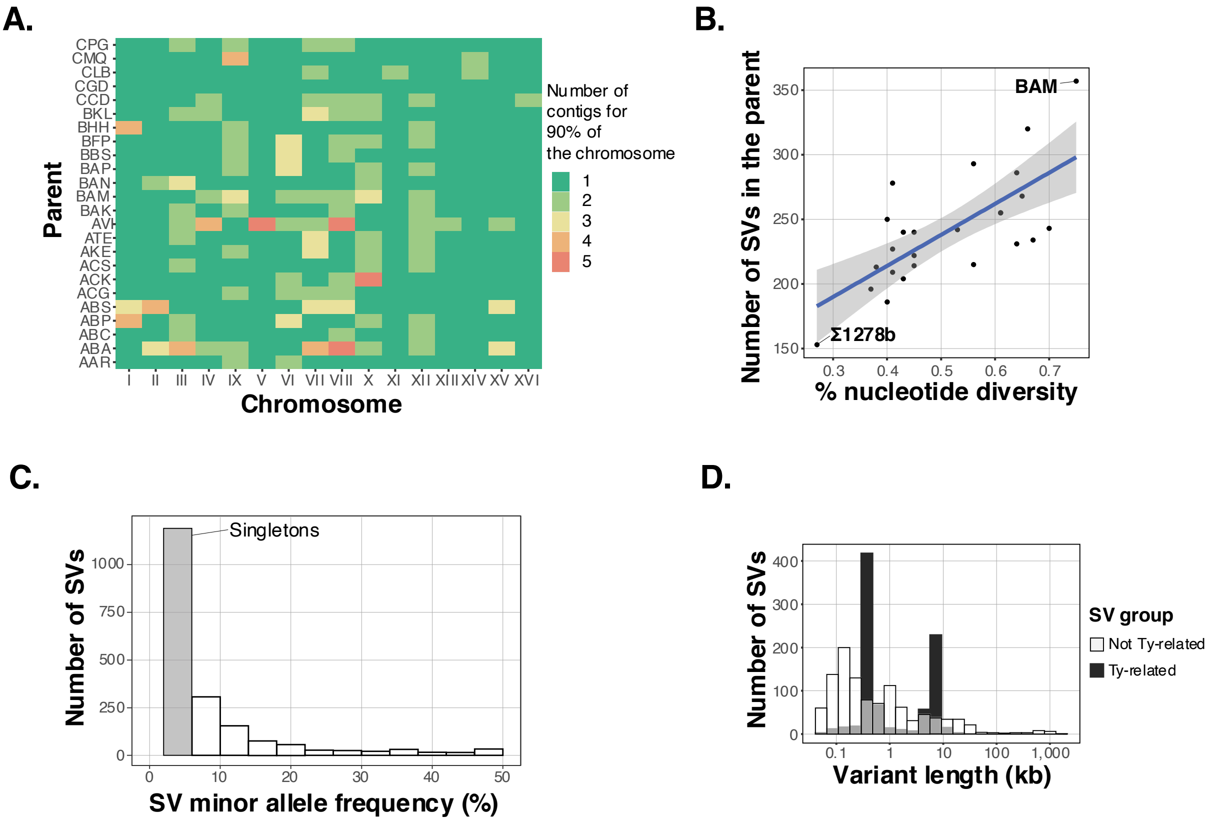


**Figure S3. Genome assemblies and structural variant detection statistics**

(A) Number of contigs, obtained by the genome assemblies of the parents, needed to cover more than 90% of each chromosome's length. (B) Correlation between the number of SVs detected in each parent and the parents nucleotide diversity from the reference strain (S288c) (correlation $r^2$=0.7, p-value=7.3e$^{-5}$). The blue line represents the linear regression line between the two variables. (C) Distribution of the SVs' MAF across the 26 parental isolates. The grey bar highlights the SVs present in only one individuals (singletons) (D) Comparison of the variant length distributions of the Ty-related SVs (black bars) and not Ty-related SVs (white bars).

### Figure S4

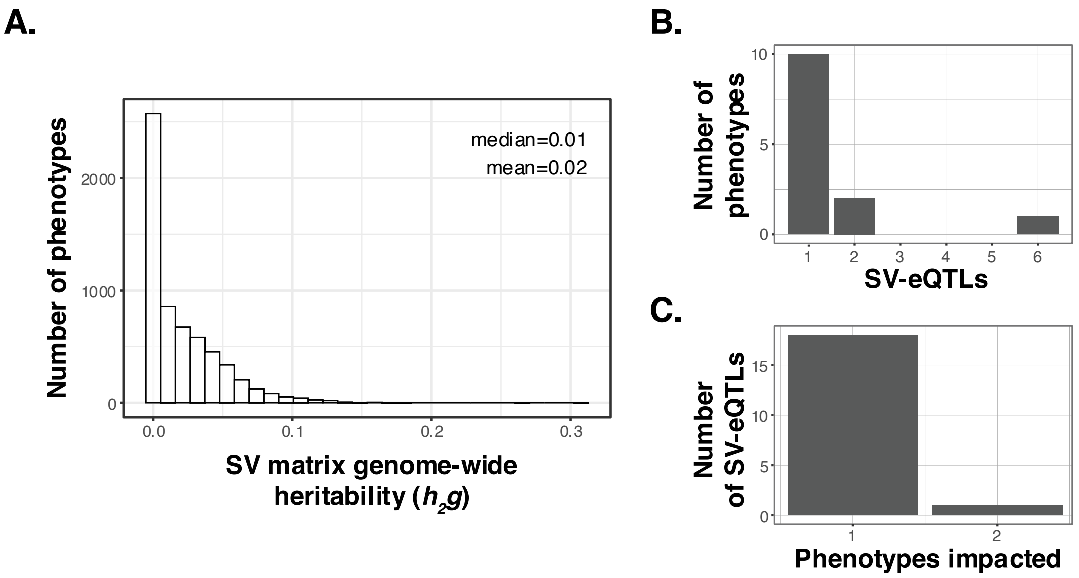


**Figure S4. SV genome-wide heritability and GWAS results**

(A) Distribution of genome-wide heritability (*$h^2g$*) based on the SV genotype matrix, excluding the SVs present in only one of the parents. (B) Distribution of the number of eQTLs associated to each phenotype. (C) Distribution of the number of phenotypes associated to each eQTL.

**Description of the supplemental tables**

**Table S1.** List of parental isolates

**Table S2.** Pairwise nucleotide diversity between the parental isolates

**Table S3.** Accessory and core genes included in the analyses

**Table S4.** Genome-wide heritability values

**Table S5.** SNP-GWAS eQTLs

**Table S6.** Nanopore sequencing metrics for the parental isolates

**Table S7.** Parental genome assembly metrics

**Table S8.** Structural variants detected

**Table S9.** SV-GWAS eQTLs

**Online data files**

**Datafile 1.** MAF of the SNPs across the diallel population and the population of 696 natural isolates.
